# Tracking Replicating HPV Genomes in Proliferating Keratinocytes

**DOI:** 10.1101/2025.03.18.644043

**Authors:** Jonathan R. Shin, Ivan Avilov, Mario Schelhaas, Franck Gallardo, Alison A. McBride

## Abstract

Human papillomavirus (HPV) genomes replicate and partition as minichromosomes alongside host chromatin during persistent infection. However, it is difficult to monitor genome dynamics in living cells because the small and compact genome will not easily tolerate expression cassettes. Here, we use ANCHOR™ technology to detect HPV18 genomes in living cells. We incorporated the cis element from ANCHOR^TM^ technology into the late region of the HPV18 genome and expressed the ParB-GFP protein from an HPV18-dependent replicon. The replicon contains the HPV18 replication origin and viral transcriptional enhancer element and can replicate stably in keratinocytes when complemented by the HPV18 genome. This small replicon expresses the neomycin resistance gene in both bacteria and eukaryotic cells and has minimal prokaryotic elements that could induce innate immunity. This molecular tool enables us to indirectly monitor the presence of the virus by detecting these fluorescent proteins in live cells and allows for real-time tracking of replicating HPV18-ANCH3 genomes in proliferating keratinocytes to inform on models of HPV genome maintenance, tethering, and amplification. Here, we visualize partitioning of the viral DNA in dividing cells and show that HPV18-ANCH genomes are distributed somewhat equally to daughter cells by random attachment to host chromosomes.

**Importance:** In persistent HPV infection, the viral genome is maintained at a constant copy number, replicates in synchrony with host DNA during S-phase, and is partitioned into daughter cells. The exact method by which HPVs partition to daughter cells is not well understood and the elucidation of such mechanisms may reveal relevant pharmacological targets to combat persistent HPV infection.

## Introduction

Persistent extra-chromosomal viral or plasmid DNA molecules must efficiently partition to daughter cells. For example, low copy bacterial plasmids use specialized partitioning mechanisms, while high copy plasmids partition by random diffusion without tethering to host structures (1). In most bacterial species, faithful partitioning is mediated by the parABS system (2). Viruses that cause persistent infection, and have extra-chromosomal genomes, must also have a mechanism to retain viral DNA in dividing cells. In gammaherpesviruses, 90% of Epstein-Barr virus (EBV) genomes are observed associated with host chromosomes as pairs and partition in a quasi-faithful manner, resulting in approximately equal distributions of genomes throughout a cell population (3). In contrast, Kaposi’s sarcoma herpes virus (KSHV) genomes are synthesized while tethered to host chromatin and partition somewhat randomly as clusters, leading to larger numbers of genomes in fewer cells of a population (4). A visual depiction of different plasmid partitioning models that could potentially represent HPV genomes can be seen in **Figure 1**.

**Figure 1.**
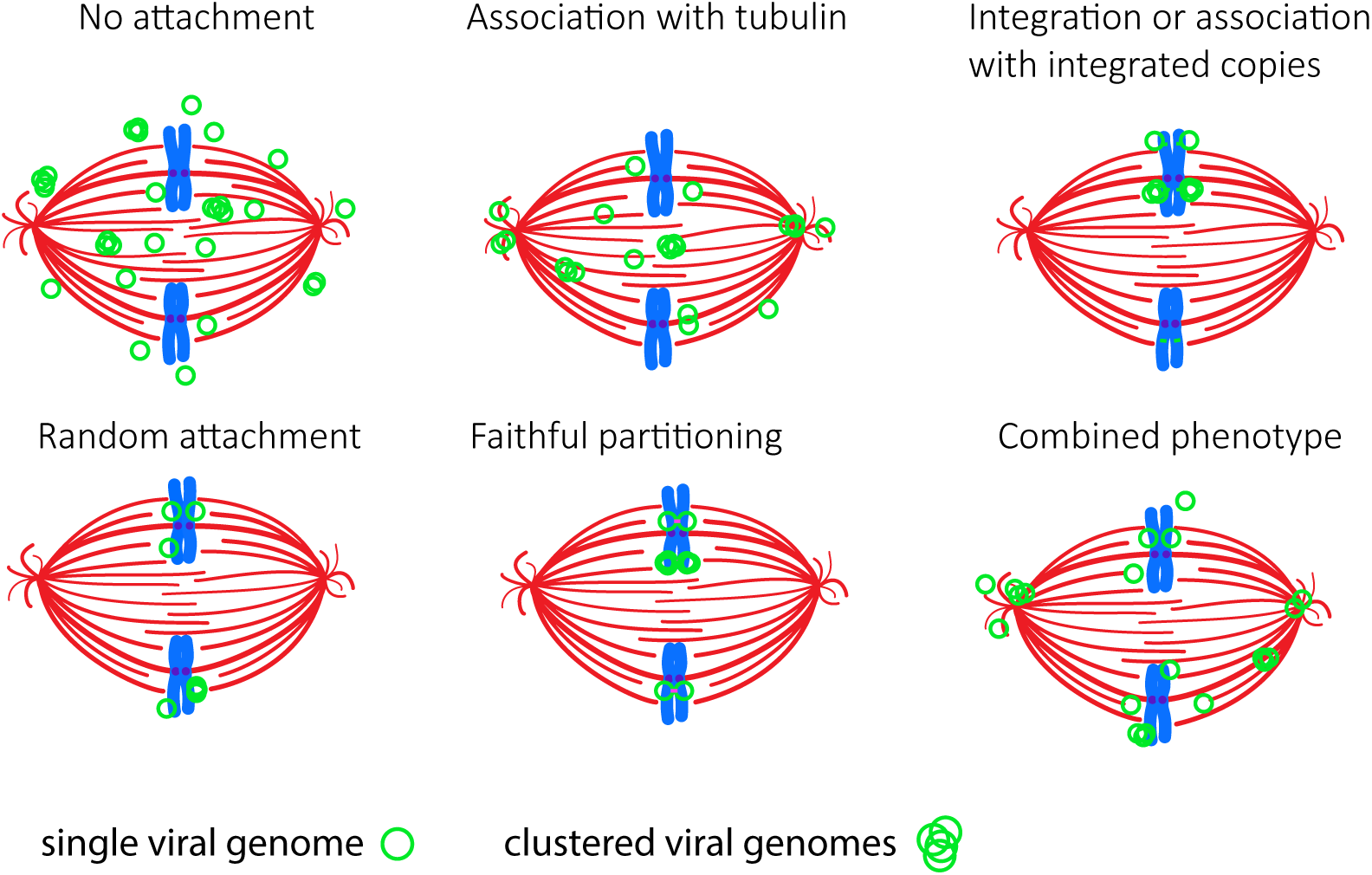
Potential HPV partitioning models. The various models show viral genomes randomly attached to chromosomes, genomes associated with tubulin, no attachment to host structures, association of viral episomes with integrated HPV genomes, or a combination of the models. Each model shows viral genomes or extra-chromosomal plasmids (*green circles*) attached pairwise to sister chromatids (*singly or in clusters)*. The pink bar represents a topological or protein link between daughter molecules.

Like other viruses with extra-chromosomal genomes, papillomaviruses are thought to maintain and partition their genomes by tethering them to host chromosomes (5–8). However, the precise mechanism of tethering is still not completely understood; in general, it is thought that the E2 protein mediates genome tethering by binding to viral DNA through its DNA binding domain and associating with host chromatin through its transactivation domain (8). Both the HPV replication origin and enhancer element in the upstream regulatory region are necessary and sufficient for efficient maintenance and partitioning of HPV-derived replicons (9). However, E2 proteins from different genera of papillomaviruses have different affinities and target regions on mitotic chromosomes (10). There has also been ongoing debate as to whether HPV genomes are “licensed” during persistent infection and thus only replicate once per cell cycle, or whether they replicate by a “random choice mechanism” whereby some genomes are amplified during S-phase and others remain unreplicated (11). Hoffman and colleagues demonstrated that HPV genomes could replicate by either mode, with increased levels of the E1 protein promoting genome amplification (11). A recent study by the Sapp laboratory showed that in some cells only a portion of the HPV genomes are associated with cellular DNA and unattached genomes could be lost to the cytoplasm necessitating reamplification in S-phase to maintain copy number (12).

Another consideration is whether each viral genome is replicated and partitioned faithfully (with the progeny genomes distributed evenly to each daughter cell) or whether chromosomal attachment and segregation is random. Both modes have been observed for the gammaherpesviruses, Epstein Barr virus (EBV) and KSHV (Kaposi’s sarcoma associated herpesvirus); EBV genomes can be observed attached to each sister chromatid and to partition in a quasi-faithful manner (3), whereas KSHV genomes form clusters that are partitioned randomly (4, 13). The HPV tracking system described here allows us to observe HPV genome partitioning in single cells in real time.

For many viruses, detection of viral DNA in live cells can be achieved by generating recombinant viral genomes that express fluorescent proteins. However, these expression cassettes can be detrimental to replication of papillomavirus genomes (14), and this approach only monitors the presence and not the location of viral DNA. Here, were able to track HPV18 genomes in real time using the ANCHOR^TM^ DNA labeling system. We inserted the ANCH3 region into the late region of the HPV18 genome and used an HPV18 derived replicon to express an OR (ParB)-GFP fluorescent protein. These minimal replicons contain a neomycin resistance gene and the HPV18 upstream regulatory region (URR) (15) and their replication and persistence relies on the synthesis of HPV18 viral replication proteins, E1 and E2. Thus, fluorescence is strictly detected in the presence of viral genomes. These URR-replicons persist in keratinocytes for many population doublings. When transfected into primary keratinocytes, both molecules replicate extra-chromosomally, and allow us to monitor the presence and location of viral genomes in dividing keratinocytes.

## Results

### Detection of HPV18 genomes using ANCHOR™ technology

NeoVirTech have developed ANCHOR™ technology to directly track DNA molecules in living cells. A cis encoded, bacterially derived partitioning sequence (ANCH3) is incorporated in the DNA molecule of interest and is specifically bound by the corresponding ParB-fluorescent protein (ParB-GFP). Initial attempts to incorporate both the cis-acting ANCH3 sequence and the gene for the trans-acting protein resulted in a replication-defective HPV18 genome. Therefore, we inserted the ANCH3 sequence alone into the late region of the HPV18 genome and expressed the ParB-GFP protein separately from an HPV18-derived replicon (**Figure 2**). ANCH3 is a specific chromosome partitioning sequence that was originally amplified directly by polymerase chain reaction (PCR) from the genome of an undisclosed bacterium (16). We inserted the ANCH3 element in both orientations in the late region of HPV18 and electroporated either wild type HPV18 or HPV18-ANCH recircularized viral genomes into keratinocytes along with a non-replicating pCGNeo plasmid. Cells were transiently selected with G418, and colonies were pooled into a cell line. The HPV18-ANCH genomes resulted in a reduced number of colonies compared to wild type, suggesting a decrease in genome establishment, but the resulting colonies still grew into established cell lines. Genomic DNA was isolated after four to six passes and viral genome replication analyzed by Southern blot. Both wild type and HPV18-ANCH genomes replicated robustly and were present at several hundred copies of extra-chromosomal genomes per cell (**Figure 2**).

**Figure 2.**
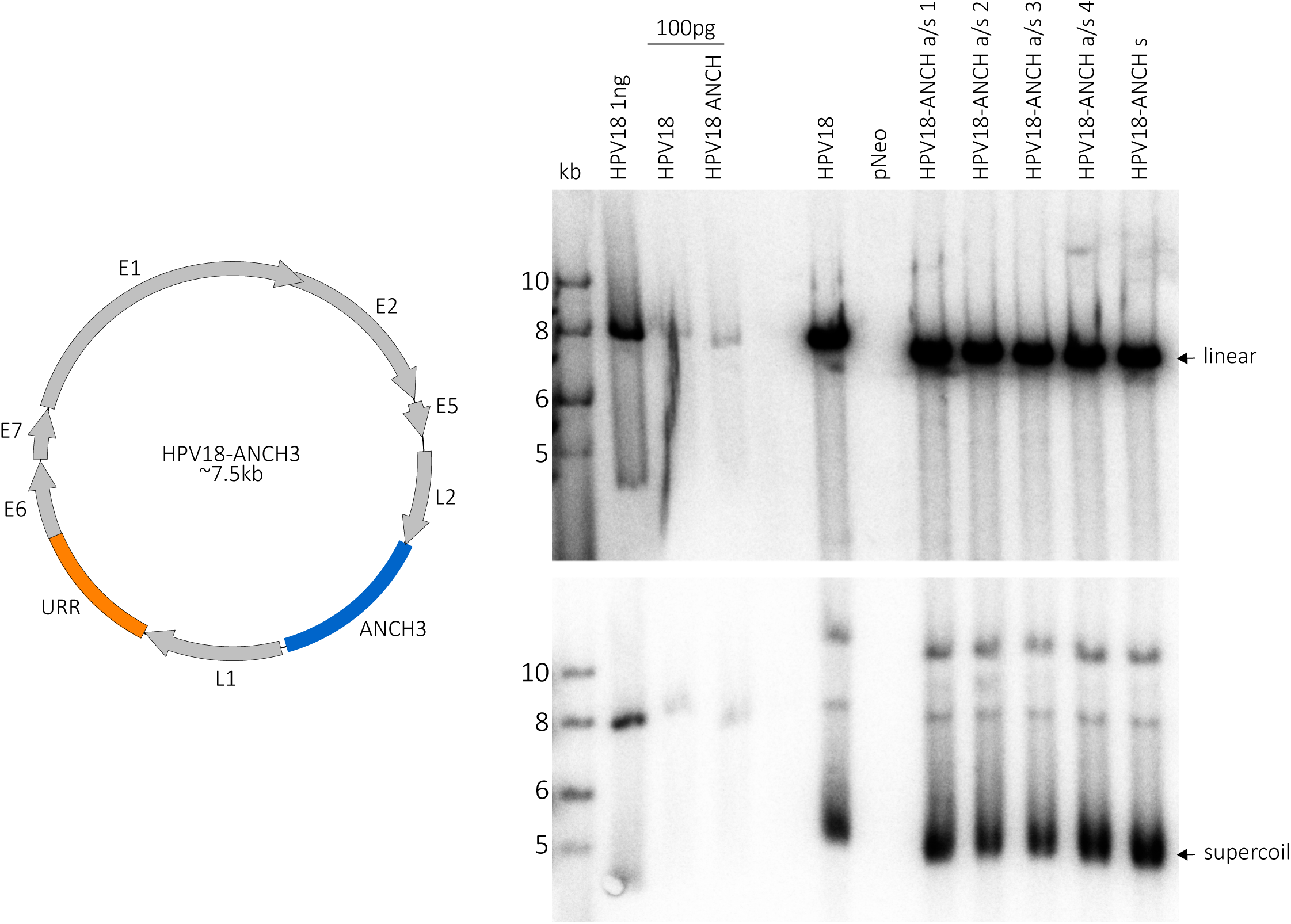
HPV18-ANCH genomes establish and replicate in keratinocytes. The ANCH3 sequence was inserted in both orientations into the late region of the HPV18 genome. HFKs were electroporated with either recircularized wild-type HPV18 or different clones of HPV18-ANCH3 genomes (s, sense; a/s, antisense) along with a neomycin resistance non-replicating plasmid, pCGNeo (pNeo), or pCGNeo_del_ (promoter deleted derivative). Genomic DNA was extracted after four to six passages and analyzed by Southern blot analysis. The top panel contains cellular DNA cleaved with AflII (which linearizes both the HPV18 and HPV18 ANCH genomes). The cellular DNA in the bottom panel was cleaved with XhoI, which does not cut either genome. HPV18 and all HPV18-ANCH3 genomes were maintained at several hundreds of copies per cell.

### Adaptation of an HPV18-derived replicon as an expression vector

We have previously shown that an HPV18 derived plasmid replicon can persist in keratinocytes containing HPV18 genomes (15). The HPV18 upstream regulatory region (URR) includes the viral replication origin with E1 and E2 binding sites for plasmid replication, partitioning, and long-term maintenance. The URR-replicon expresses the neomycin resistance gene in both bacteria (I-ECK1 promoter) and mammalian cells (SV40 promoter) and has minimal prokaryotic elements and an overall reduction of CpG dinucleotides to avoid foreign DNA defenses such as TLR9 detection and methylation of CpG residues. Here, we adapted the replicon to serve as a long-term expression vector in keratinocytes in the presence of the HPV18 genome. We identified four insertion sites, between functional elements in the replicon, and inserted a short multiple cloning site polylinker in both orientations resulting in eight different URR-replicon vectors. Each plasmid was electroporated into human foreskin keratinocytes (HFKs), or HFK-HPV18 cells to assess its ability to establish as a stable URR-replicon. All eight URR-replicons gave rise to neomycin-resistant colonies and as expected, only in the presence of HPV18 (**Supplemental figure 1).**

### Assessment of long term GFP expression from different positions within the URR replicon

Initially, to test the ability of the replicon to serve as an expression vector in keratinocytes, we used a gene encoding maxGFP, an enhanced GFP derived from the copepod *pontellina plumata*. In the original pmaxGFP plasmid, GFP is expressed from a powerful CMV promoter and uses an SV40 polyadenylation site. Although the CMV promoter gives robust expression, we have found that such high levels of GFP are not optimal for sustained, long-term gene expression and cell health. Therefore, we cloned the GFP gene (along with a chimeric intron, which likely reduces HUSH-mediated repression of intronless DNA (17), into pSelect-puro which uses a hybrid hEF1/HTLV promoter and SV40 polyadenylation site to express a gene of interest. In other studies in our laboratory, we have found this promoter gives sustained long-term expression in keratinocytes. Both plasmids were transiently transfected into HFKs that were either conditionally immortalized by the ROCK inhibitor Y27632 (SA cells), or by HPV18 (18). As shown in Supplementary Figure 2, transferring the intron and GFP gene into pSelect-puro reduced GFP expression to a more moderate level in both cell lines. Additionally, the GFP signal was greatly increased in the HPV18 positive cells. We have not investigated this enhancement further but surmise that it is due to HPV18 mediated inhibition of innate immunity to foreign DNA (e.g. cGAS/STING). We view this effect of HPV18 as beneficial to our project as it increases the establishment of URR-replicons in keratinocytes.

We inserted the optimized hEF1/HTLV pmaxGFP expression cassette into each of the eight polylinker URR-replicons and electroporated them into keratinocytes containing HPV18 genomes. Of note, there was a substantial reduction in drug resistant colonies compared to the parental URR-replicon but, although establishment was reduced, each HPV18 cell line eventually grew out under G418 selection. In parallel, DNA was collected six days post-electroporation and analyzed by Southern blot using a probe against either the HPV18 genome or the replicon to determine whether the expression cassette hampered viral replication. As shown in **Supplemental Figure 3**, some of the pmaxGFP URR-replicons replicated transiently but with much less efficiency than the empty URR-replicon. Endogenous HPV18 genomes were maintained at high copy number in all cells **(Supplemental Figure 3B**). After continued passage under G418 selection (∼4 weeks or three passages), DNA was prepared to determine whether the replicons were stably maintained. Of note, GFP was expressed to the degree that cell pellets were fluorescent green and individual cells expressing GFP appeared unhealthy by microscopy. We concluded that the pmaxGFP protein itself was toxic and the full length pmaxGFP-URR-replicon was not well tolerated by the cells. Because of this toxicity, we decided not to proceed with the pmaxGFP gene and to focus on expression of the ParB-GFP protein that could bind and track HPV genomes.

### Generation of a ParB-GFP Expression Replicon

The ParB-GFP protein is a fusion between the bacterial ParB protein and a synthetic, monomeric non-Aequorea fluorescent protein named Dasher (ATUM). To determine whether expression of this protein was better tolerated by keratinocytes, we first inserted the ParB-GFP open reading frame into the pSelect vector between the hEF1/HTLV promoter and the SV40 polyadenylation site, and then transferred the expression cassette into the NheI-sense position of the URR-replicon.

The parental URR-replicon, and the ParB-GFP URR-replicon were electroporated into cells containing either HPV18 or HPV18-ANCH genomes (cells described in Figure 2) and cells were either selected with G418 for 14 days, or genomic DNA collected after six days without selection for transient replication analysis. As shown in Figure 3, the ParB-GFP replicons gave rise to colonies in HPV18 containing cells, but the number and size of colonies were greatly reduced compared to the parental URR-replicon. This is similar to the observations made with the pmaxGFP replicons, indicating that replicons encoding expression cassettes are established at much lower frequency than the empty URR replicon.

**Figure 3.**
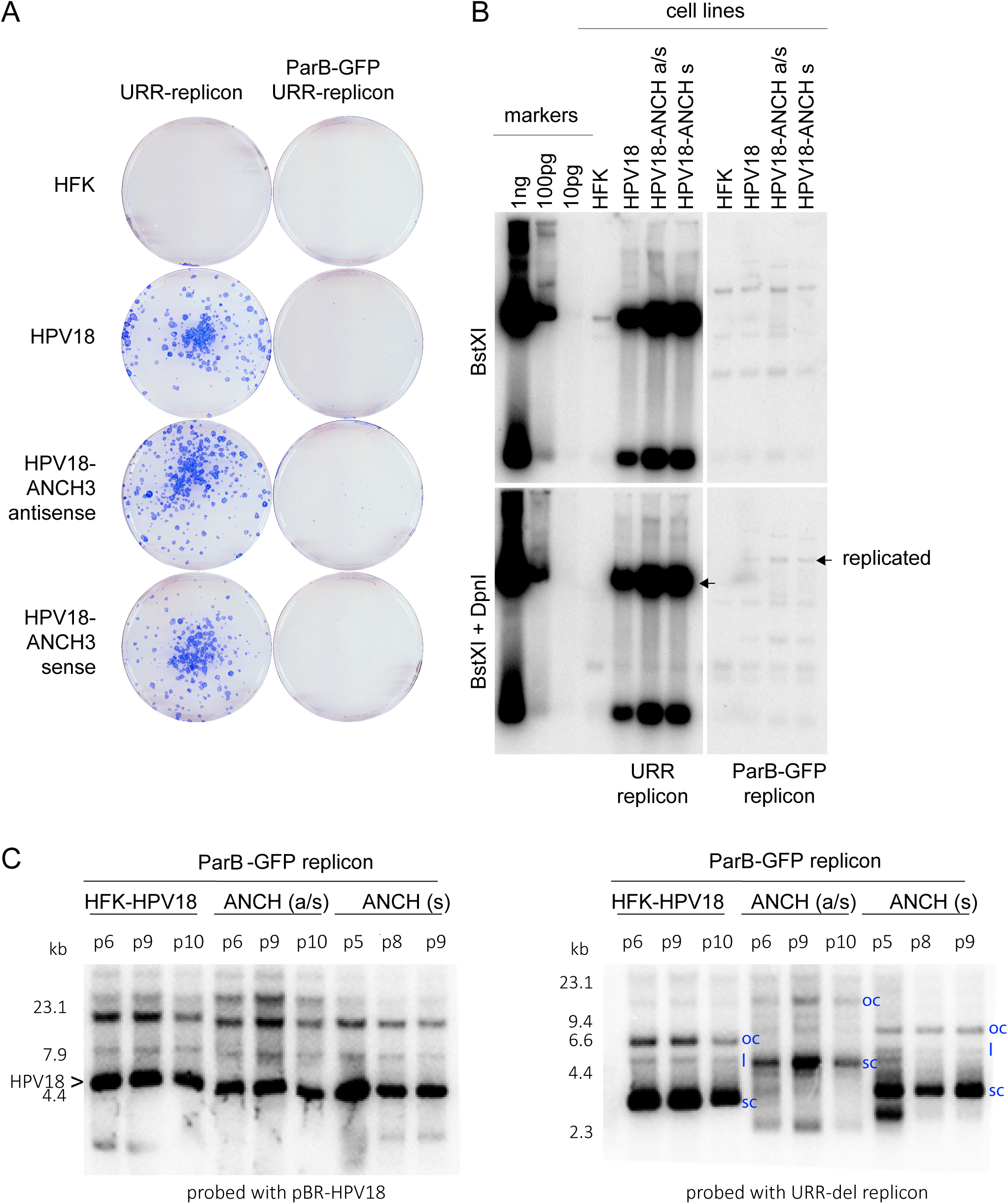
Establishment and replication of the ParB-GFP-URR-replicon in HFK, HPV18 and HPV18-ANCH cell lines. (A) The ParB-GFP-URR-replicon was electroporated into the cells indicated and colonies were stained after 14 days under G418 selection. (B) DNA was harvested from a parallel set of plates six days post-electroporation without G418 selection. The cellular DNA in the top panel was digested with BstXI and in the bottom panel with BstXI and DpnI to detect DpnI resistant replicated DNA. Size/copy number markers are loaded on the left. A longer exposure is shown for the samples containing the ParB-GFP replicon because of differences in replication efficiency. (s, sense; a/s, antisense). (C) The three cell lines shown in panel B were cultured for the number of passes shown at the top of each lane. DNA was prepared from each cell line, digested with PacI (a non-cutter for both HPV18 and the ParB-GFP replicon) and probed with the plasmids indicated along the bottom. The positions of supercoil (sc), open circle (oc) and linear (l) replicons are indicated in each cell line.

For transient replication analysis of the ParB-GFP replicons, genomic DNA samples were digested with BstXI and DpnI and subjected to Southern blot analysis. As shown in **Figure 3B**, the ParB-GFP expression replicon did replicate, but at greatly reduced levels compared to the parental URR-replicon. Nevertheless, the colonies containing the ParB-GFP expression replicon grew into an established cell line. Southern blot analysis of cellular DNA collected between passes p5 and p10 showed that the replicons (and HPV18 ANCH genomes) stably replicated over time although the URR-replicons in the HPV18-ANCH (a/s) cells had likely undergone multimerization (Figure 3C).

### Location of the ParB-GFP protein in HPV18 and HPV18-ANCH cell lines

In parallel, cells were screened and analyzed by live cell fluorescence microscopy multiple times during the establishment of the pooled HPV18-ANCH ParB-GFP URR-replicon cell lines. This showed strong but variable cytoplasmic GFP fluorescence in many cells containing the ParB-GFP URR-replicon. There were multiple phenotypes including diffuse ParB-GFP signal throughout the nucleus, and various levels of ParB-GFP intensity. However, cells containing the HPV18-ANCH genome routinely contained punctate ParB-GFP signals in the nucleus, while HPV18 containing cells never did. **Figure 4** shows an example of two such colonies captured in live cells after more than 33 days of G418 selection. The ParB-GFP signals were monitored along-side DNA (SPY-650) and tubulin (SPY-555) live-cell dyes.

**Figure 4.**
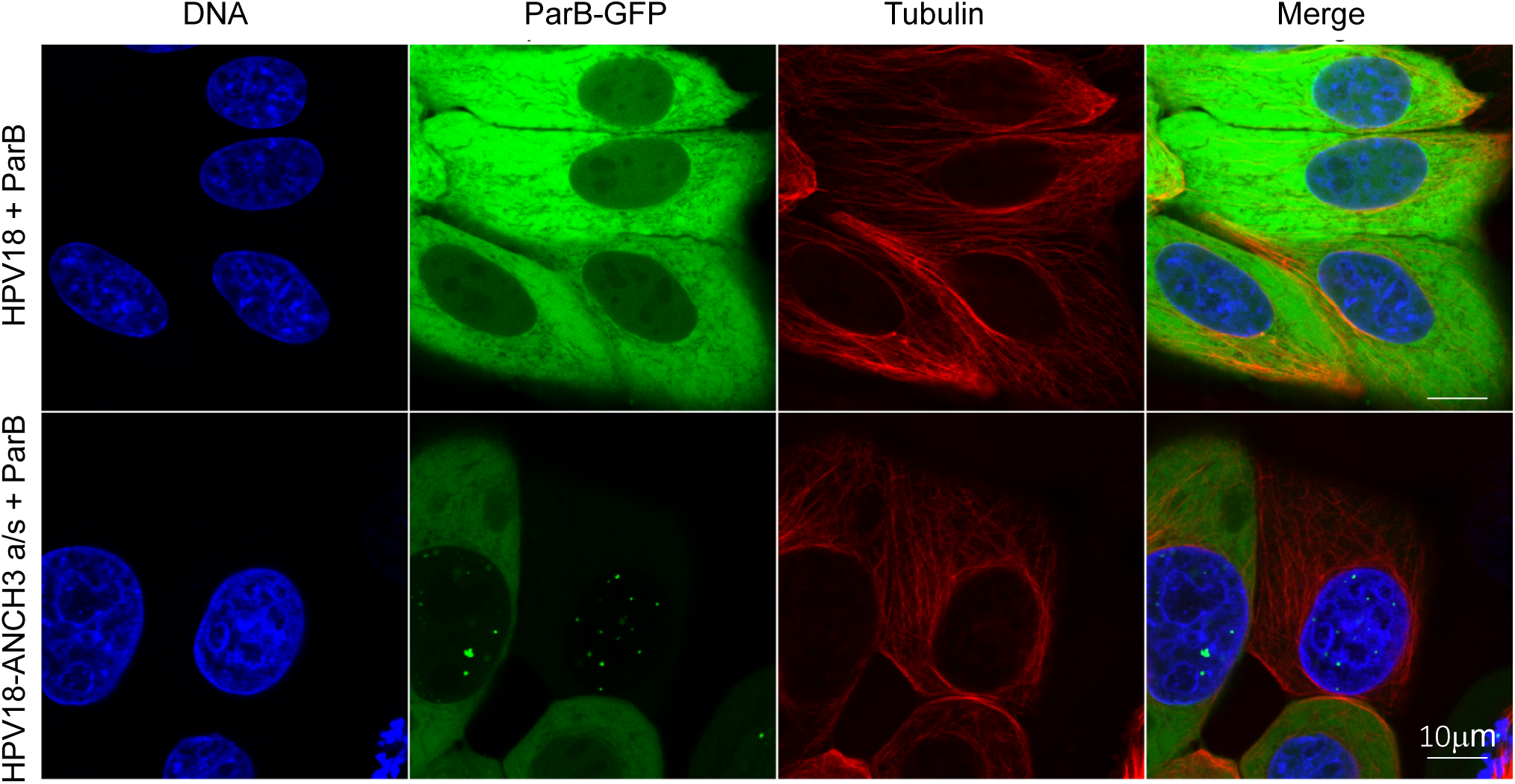
Detection of HPV18-ANCH3 genomes using ANCHOR^TM^ technology. Shown are confocal microscope image slices of ParB-GFP fluorescence in live HFKs after over 33 days of G418 selection. The top row contains wild-type HPV18, and the bottom contains the recombinant HPV18-ANCH genome. Cells with the HPV18-ANCH genome contain punctate ParB-GFP signals localized in the nucleus. Scale bar is 25 µm. Cells were incubated with live-cell DNA (SPY-650) and tubulin (SPY-555) dyes. Scale bar is 10 µm

To ensure the punctate ParB-GFP signals represented HPV18-ANCH genomes, cells were plated on coverslips and HPV18 DNA was detected by FISH, and the ParB-GFP by a specific antibody to DasherGFP. The antibody was required to detect the position of ParB-GFB because the FISH protocol destroys its fluorescence. As shown in **Figure 5**, the ParB-GFP signals co-localize with nuclear HPV18-FISH foci and the most prominent puncta are indicated with white arrows.

**Figure 5.**
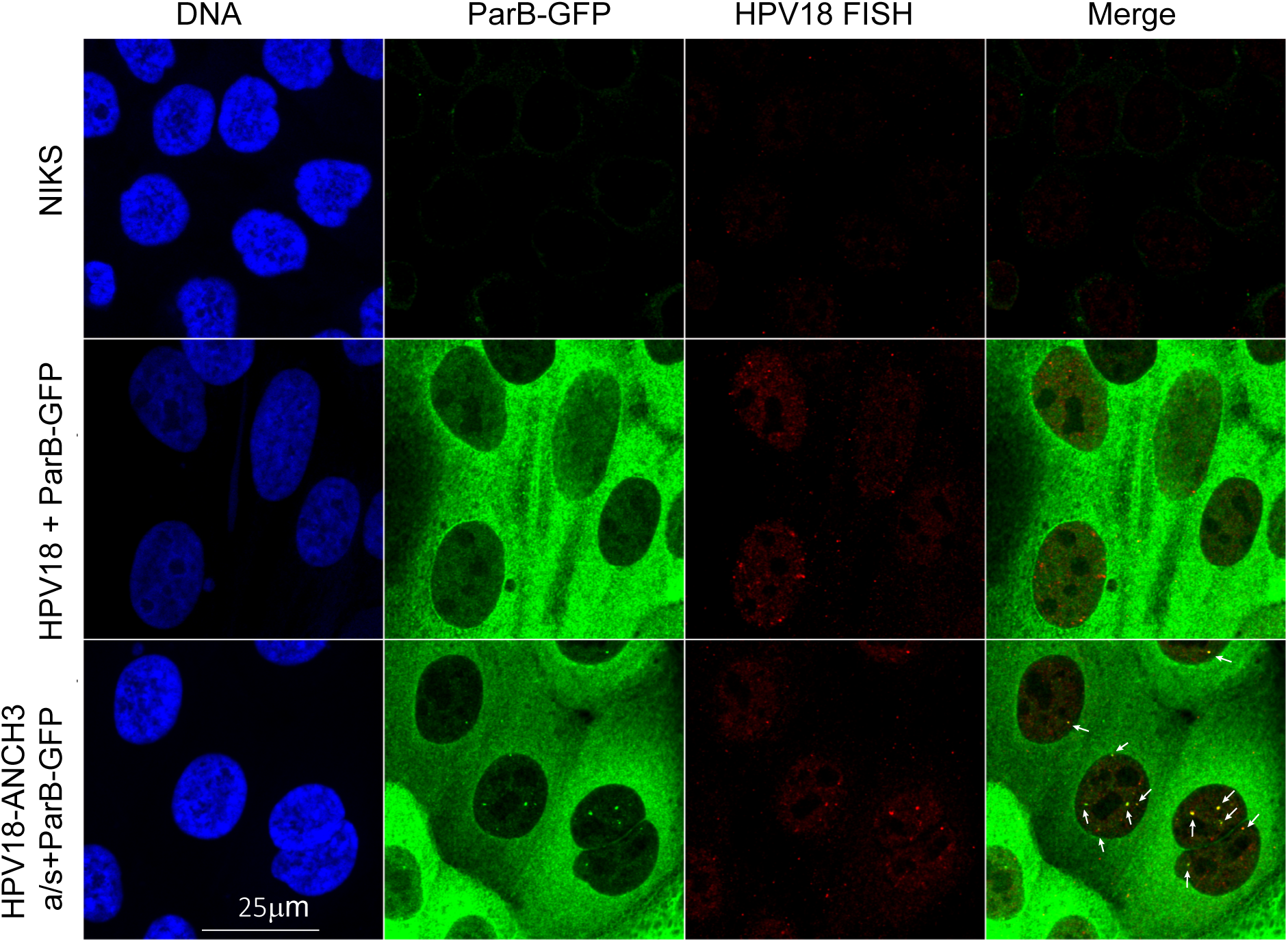
ParB-GFP puncta co-localize with HPV18-ANCH3 FISH signals. Shown are combined immunofluorescence images using an anti-DasherGFP antibody (CometGFP, ATUM 02) and a URR-negative HPV18 DNA as a FISH probe to detect viral genomes. Normal Immortalized Keratinocytes (NIKS) were used as a negative control for viral genomes and ParB-GFP expression. White arrows identify ParB-GFP puncta.

Additionally, single cell clones were isolated from the original HPV18-ANCH pools to observe uniform phenotypes during fluorescence microscopy. Cell clones varied in general size, rate of growth, and ParB-GFP expression and HPV18 copy number. Three cell lines were used for detailed live cell microscopy and their characteristics are summarized in **Table 1**. (1) HFK HPV18 ParB-GFP cells contained the wildtype HPV18 genome and expressed ParB-GFP in a uniform cytoplasmic fashion (2) HFK-HPV18-ANCH ParB-GFP-clone contains the ParB-GFP replicon and has a low copy number of the HPV18-ANCH genome that is observed as one to four nuclear foci in most cells. (3) HFK HPV18-ANCH a/s ParB-GFP is a non-clonal cell line containing the HPV18-ANCH genome and the ParB-GFP replicon, and contains multiple and heterogeneous foci of ParB-GFP in many cells.

**Table 1.**

| Cell Line | Abbreviation | Viral genome | Replicon | ParB-GFP fluorescence phenotype | Viral genome copy number | Clonal | Used For: |
| --- | --- | --- | --- | --- | --- | --- | --- |
| NIKS | NIKS | None | None | None | - | No | Negative control for viral genomes and GFP |
| HFK | HFK | None | None | None | - | No | Negative control for viral genomes and GFP |
| HFK HPV18 | HPV18 | HPV18 | None | None | High | No | Parental cell line, negative control for GFP |
| HFK HPV18-ANCH | HPV18-ANCH | HPV18 | None | None | High | No | Parental cell line, negative control for GFP |
| HFK HPV18 ParB-GFP | HPV18 + ParB | HPV18 | ParB-GFP | All cells have diffuse cytoplasmic signal | Low | Yes | Negative control for ANCH3 viral genomes |
| HFK HPV18-ANCH a/s ParB-GFP-pool | HPV18-ANCH a/s + ParB | HPV18 ANCH-antisense | ParB-GFP | Mixed phenotype: GFP negative; diffuse cytoplasmic signal; diffuse nuclear signal; cells with nuclear puncta and/or cytoplasmic puncta | High | No | Tracking HPV genomes |
| HFK-HPV18 - ANCH ParB-GFP-clone | HPV18-ANCH + ParB | HPV18 ANCH-sense | ParB-GFP | All cells have diffuse cytoplasmic signal and small nuclear puncta | Low | Yes | Observing E1/E2-induced HPV18-ANCH replication factories labeled by ParB-GFP |

### Analysis of HPV18 Genome Partitioning in Live, Dividing Cells

One notable feature of HPV genome replication is the ability of viral genomes to persist extra-chromosomally at constant copy number, for many cell generations. There has been much debate about whether viral genomes are licensed (replicate once per cell cycle) or replicate by random choice mechanisms (8). Raj and colleagues showed that it depended on both viral type and individual cell line (11). There is compelling evidence that some papillomavirus genomes (e.g. BPV1) are associated with mitotic chromosomes (6, 7), but less evidence for HPV genomes. A recent study showed that the viral genome copy number dramatically decreased in mitosis and was reamplified in the following S-phase (12).

To examine maintenance and partitioning of the HPV18-ANCH genomes, we imaged the HPV18-ANCH cell lines expressing ParB-GFP from the URR-replicon on chambered coverslips at 37°C, 5% CO_2_, by confocal microscopy for periods of 12 to72hrs, with or without cell-permeable live-cell stains for DNA and tubulin. In general, the HPV18-ANCH genomes were observed associated with mitotic chromosomes, though unattached genomes could also sometimes be observed. In metaphase, HPV18-ANCH genomes could be observed associated with host chromosomes aligned along the metaphase plate and then partitioned on chromosomes in anaphase and telophase. However, in some cases a portion of the viral DNA/ParB-GFP signal was left behind, still aligned along the metaphase plate. Furthermore, viral DNA (or ParB-GFP signal) that was not attached to host chromosomes could be observed lost to the cytoplasm when the nuclear membranes reformed.

**Figure 6** shows examples of these different phenotypes in time lapse images of five mitotic cells. Cell One shows an example of HPV genomes/ParB-GFP puncta bound to mitotic chromosomes and distributed evenly to daughter cells, while in Cell 4 the HPV genomes are partitioned unevenly. We often observed ParB-GFP signals moving into the cytoplasm after mitosis (e.g. cell 4, 5). As described above, ParB-GFP puncta (likely extra-chromosomal) often align with metaphase chromosomes. However, some puncta do not stay co-localized with DNA during anaphase and are left behind in the cytoplasm. There are also instances of single ParB-GFP foci that duplicate and faithfully partition upon anaphase, and these could represent viral genomes integrated into host chromosomes, or a cluster of genomes (e.g. cell 3).

**Figure 6.**
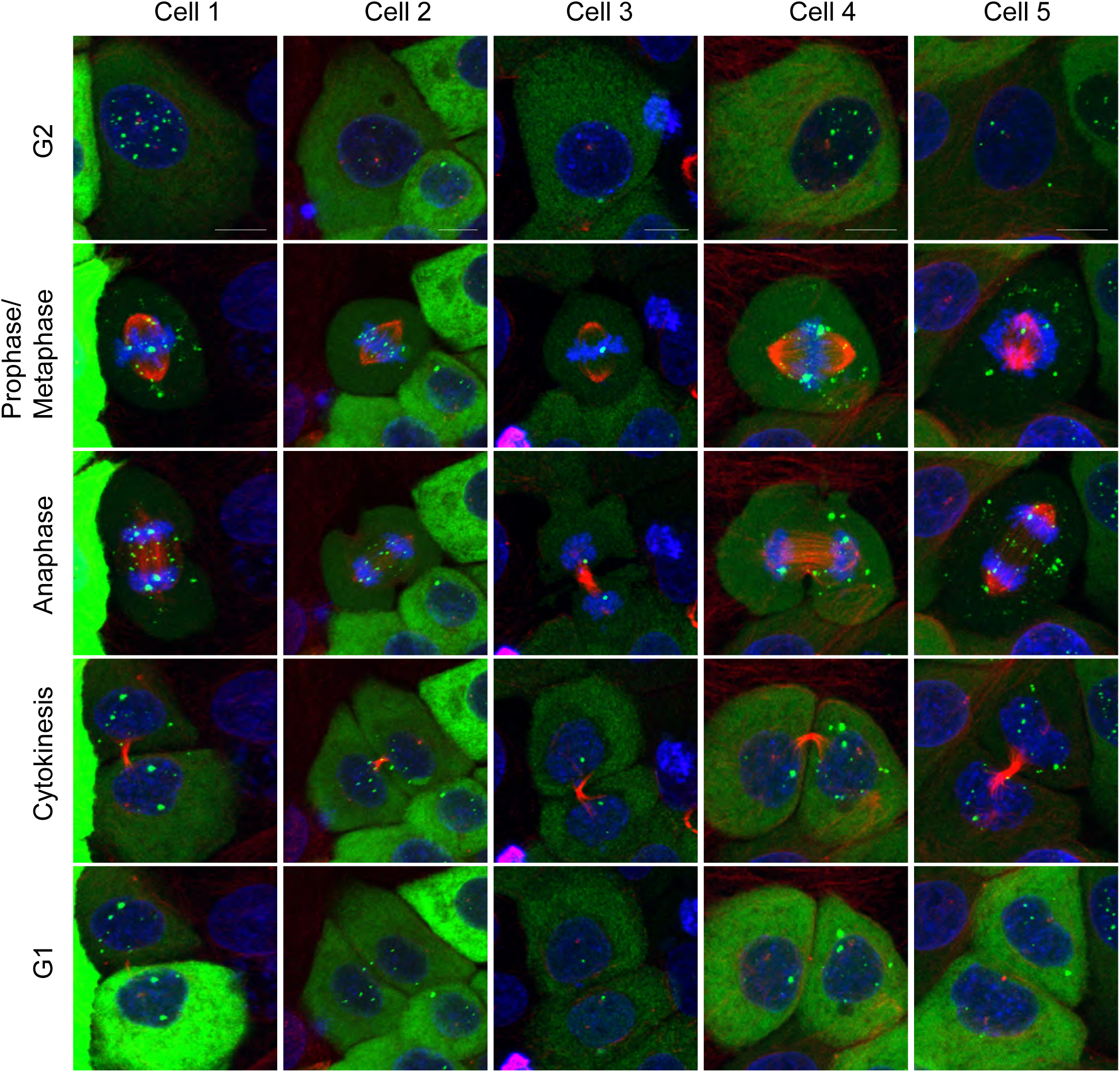
Partitioning of ParB-GFP puncta. These live confocal time lapse images capture dividing HFKs harboring replicating HPV18-ANCH genomes. The ParB-GFP protein (*green*) is expressed from the URR-replicon. Additionally, DNA (*blue*) and tubulin (*red*) dyes are present. Distinct ParB-GFP dots are seen associating with host chromosomes or failing to associate with chromosomes. These distinct signals can duplicate in number during mitosis and are partitioned equally or unequally between daughter cells (scale bar is 15µm).

**Figure 7** show the distribution of partitioning phenotypes observed in 26 different HPV18-ANCH a/s + ParB mitotic cells from several live-cell time lapse experiments. Cells were categorized in several different ways. ParB signals were observed in puncta (dots) in the mitotic cells, and these were observed either exclusively associated with mitotic chromosomes (16.7%) or localized to both mitotic chromosomes and the surrounding cytosol (75%). The ParB puncta were distributed to daughter cells and each cell was analyzed to determine whether this distribution was split evenly between the daughter cells (56%) or somewhat unevenly (44%). The mitotic cells were examined carefully to see if partitioning was faithful (a single puncta aligned on the metaphase chromosomes that splits in two as the chromosomes separate) and this was only clear in 13.4% cells. More frequently (86.7%) the partitioning appeared random with different puncta segregating to different daughter cell. Finally, in most cells (80.8%), a proportion of ParB puncta were shed to the cytoplasm as cells proceeded through mitosis. These puncta were often associated with host chromosomes in metaphase, but frequently as the chromosomes moved apart in anaphase (with bound ParB puncta) a line of ParB signal could be observed left behind along the metaphase plate (see **Figure 6** Cell 5 for an example). In summary, ParB puncta that were associated with host chromosomes partition efficiently to daughter cells, but those that were not associated were often lost to the cytoplasm after cytokinesis.

**Figure 7.**
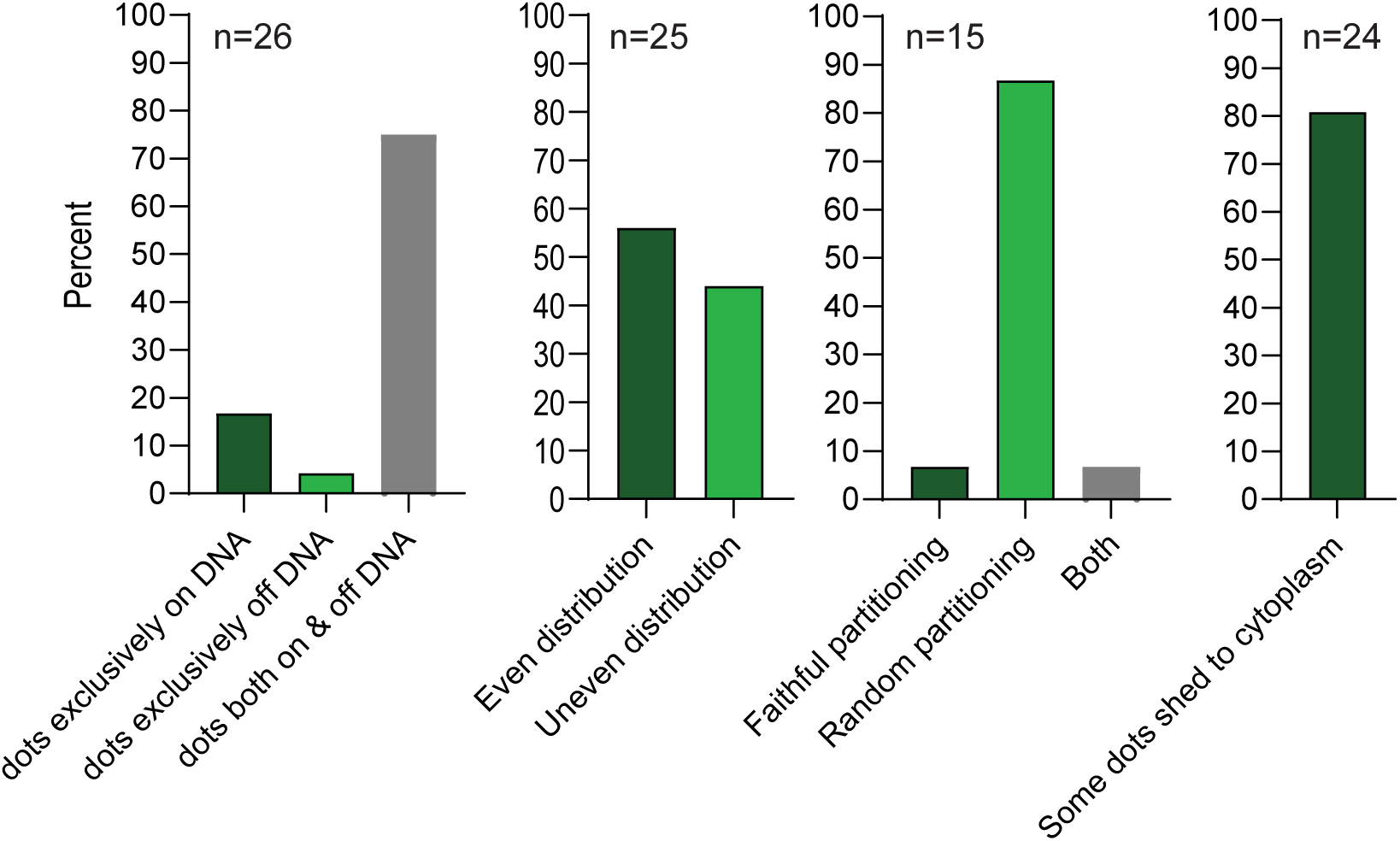
ParB-GFP puncta partitioning phenotypes. Shown are percentages of certain ParB-GFP phenotypes observed in 22-26 live cells containing the HPV18-ANCH genomes.

Six mitotic HPV18-ANCH mitotic cells were selected for detailed analysis (**Figure 8**). Live cell movies of each cell were analyzed frame by frame for parB-GFP puncta. The number of puncta at each stage of mitosis were counted and characterized as Nuclear/On chromosomes or Cytoplasmic, where possible. Note that the puncta were not of uniform size (see **Figure 6** for examples), implying that the larger dots were composed of clustered genomes. The number of nuclear puncta in the G2 parental cells (average number ∼ 14) was more than double that of the G1 daughter cells (average number ∼6) and there were more cytoplasmic puncta in the G1 daughter cells (an average of 0.67 in G2 and 2.8 cytoplasmic puncta in G2 cells) that probably originated from genomes that were not attached to host chromatin during mitosis. In some metaphase cells (Cells D, E, and F), the number of puncta increased significantly but this was due to a large proportion that were not attached to host chromosomes. In anaphase and telophase, the number of puncta were distributed quite evenly between daughter cells, consistent with the quantitation in **Figure 7**. In conclusion, HPV18-ANCH genomes are distributed to daughter cells by attachment to host mitotic chromosomes, but unattached genomes are shed to the cytoplasm and are often detected in the cytoplasm of the daughter G1 cells.

**Figure 8.**
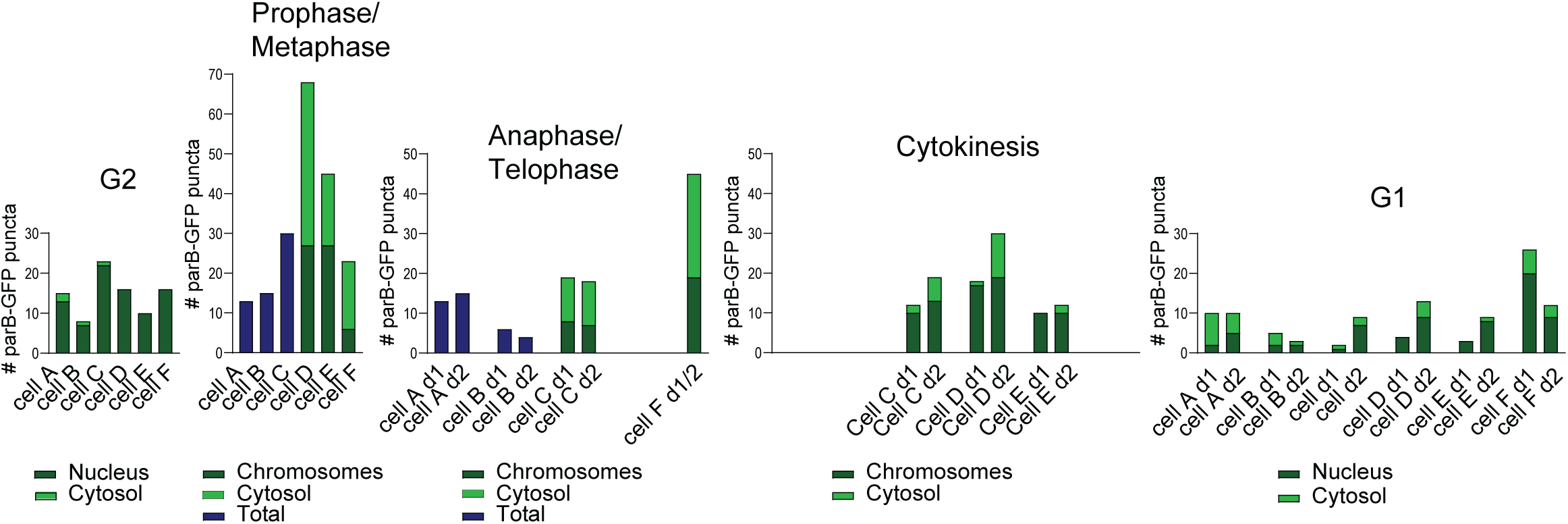
Detailed quantitation of six mitotic cells. Six mitotic cells were selected from live cell movies of HPV18-ANCH cells and analyzed frame by frame for ParB-GFP puncta. The number of puncta were characterized as Nuclear/On chromosomes or Cytoplasmic, where possible (as shown in dark green/light green, respectively.) When it was difficult to characterize this phenotype, the total number of puncta are given (dark blue bar). Note that the puncta were not of uniform size, implying that the larger dots were composed of clustered genomes (see Figure 6 for examples).

We next proceeded with automated analysis of the live cell time-lapse images, focusing on relevant mitotic stages (**Figure 9A**). First, we quantified the total number of viral genome puncta across mitosis (**Figure 9B**), by automatic spot intensity thresholding. A clear, yet variable, increase in the number of genome spots became apparent upon nuclear envelope breakdown (NEBD) and peaked at prometa/metaphase (**Figure 9B**). As this coincided with influx of ParB-GFP upon NEBD, we surmise that we detect additional genome signals that were below the background threshold prior to NEBD; increased concentration and molecular crowding effects could increase loading onto the genome. In support of this, the number of foci decreased with the formation of two daughter cells, which were quantified as a single unit. In addition, there was also an overall increase in spot intensity (not quantified). Second, to determine the association of viral genome spots with chromatin, we used intensity and shape-based image segmentation of the DNA signal to establish a three-dimensional chromatin mask. Then we classified the previously quantified viral genome puncta depending on the overlap with segmented chromatin (**Figure 9C**). A significant proportion of viral genome puncta (∼40%) dissociated from chromatin after NEBD and remained in the cytosol after cytokinesis. In addition, half of the chromatin-dissociated puncta transiently associate with the spindle (**Supplementary Figure 4**). Both findings suggest that chromatin condensation displaced a certain number of viral genomes from chromatin and/or they were never fully tethered, rather than displacement by forces generated by the spindle.

**Figure 9.**
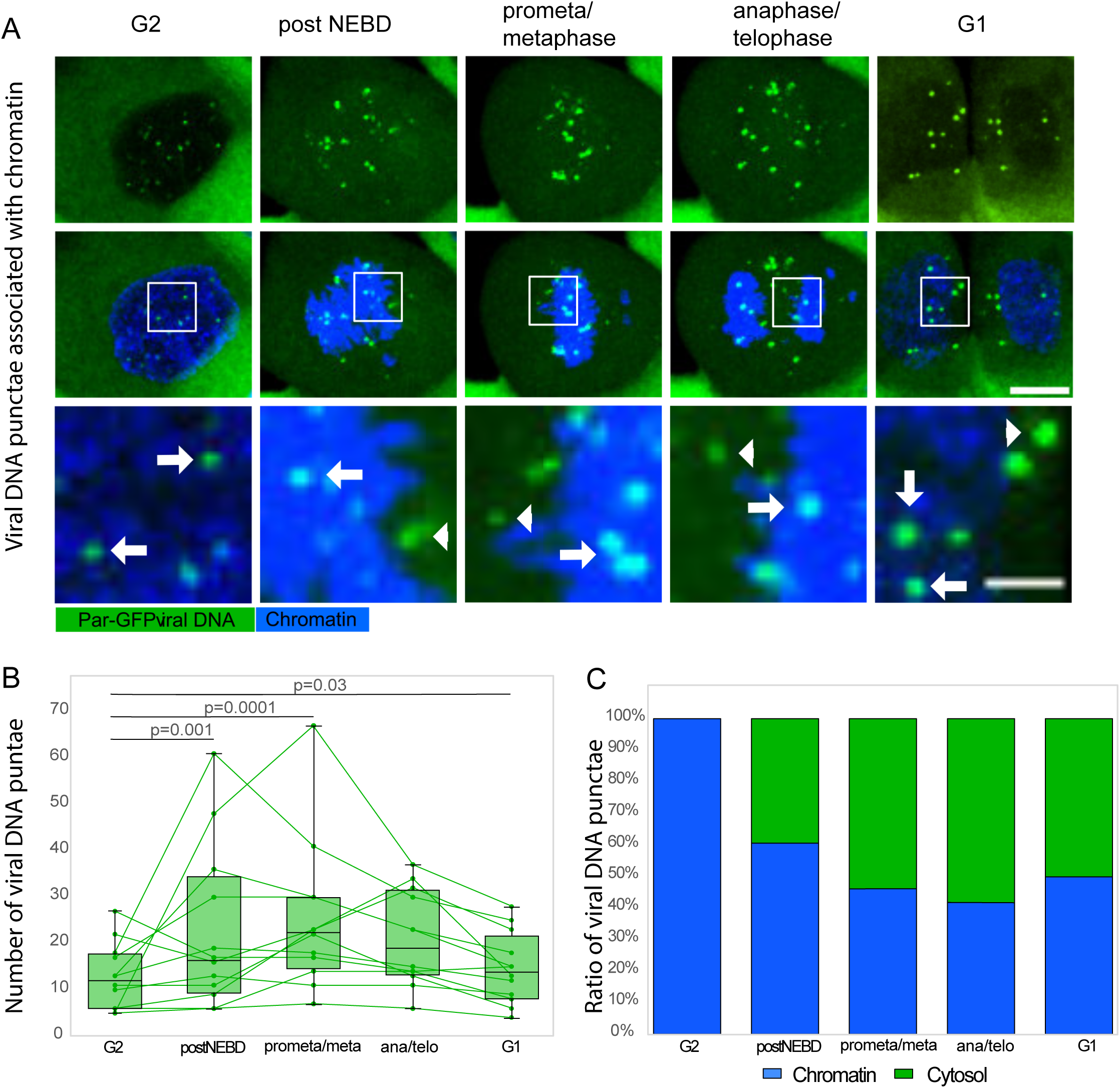
HPV18 genomes dissociate from chromatin predominantly upon nuclear envelope breakdown (NEBD) A. Representative confocal time-lapse image of HPV18-ANCH cell bearing viral genome puncta (ParB-GFP, green) such as used for subsequent image analysis. Mitotic stage classification is based on labeling for chromatin (SPY-650-DNA, blue). Solid arrows indicate viral genomes associated with chromatin (classified as ‘chromatin’, see Figure 9C), arrowheads indicate viral genomes that are not associated with chromatin (classified as ‘cytosol’, see Figure 9C). Maximum intensity projections are shown, scale bar is 10 μm, for crop-ins 3 μm. B. Boxplot showing dynamics of viral genome numbers, quantified by automated signal intensity thresholding during indicated mitotic states (see material and methods for details). Viral genome puncta significantly increase upon NEBD, peak during prometa/metaphase, and decrease to about interphase levels after mitosis. Green lines represent dynamics of viral genome numbers (green dots) for each quantified cell. N=12, the two nascent daughter cells were quantified as one unit. All statistical comparisons were done using two-tailed T-test. C. Normalized ratios between viral genomes that overlap with automatically segmented chromatin (‘chromatin’, blue, indicated in Figure 9A with arrows) and non-overlapped ones (‘cytosol’, green, indicated in Figure 9A with arrowheads) during indicated mitotic stages (see material and methods for details). A significant proportion of viral genome puncta (∼40%) already dissociate from chromatin upon NEBD. About the same number of genomes remain in the cytosol of the daughter cells.

## Discussion

Here, we present a system for tracking HPV18 genomes in live keratinocytes. There are several ways to track viral genomes; most common of which is inserting a gene encoding a fluorescent protein into the viral genome to monitor the presence of viral DNA (19, 20). However, there are also examples of cis tracking elements, such as tandem arrays of the lac operator, being inserted into viral genomes to directly detect viral DNA (4, 21). However, HPV presents more challenges due to the compact design of the viral genome, which contains many overlapping cis elements and does not tolerate many foreign DNA cassettes. In addition, HPV genomes will only replicate in keratinocytes in coculture with feeder fibroblasts, and these primary cells are often refractory to expression of foreign DNA.

Here, we use ANCHOR™ technology and inserted an ANCH3 element into the late region of HPV18 in a site known to tolerate foreign cassettes (19). We expressed the ParB-GFP ANCH-binding protein from a minimal replicon that only replicates in the presence of the HPV18 E1 and E2 replication proteins (15). This replicon has minimal bacterial elements, is CpG free, provides neomycin resistance, and persists in keratinocytes indefinitely (9). Here we modify and optimize the replicon to serve as an expression vector in HPV18 positive cells.

The ANCHOR™ system has successfully tracked many DNA viruses at different stages of infection. These include baculovirus, cytomegalovirus, equine herpesvirus, poxvirus, and adenovirus (22–26). Here, we show that the ANCHOR™ system can also track small DNA virus genomes and these genomes can be followed in long-term persistent infection. The cell lines described in this study have been passed up to ten times and still retain the HPV18-ANCH genomes and the ParB-GFP expressing replicon, and consistently express ParB-GFP.

It has been well established that HPV genomes are partitioned to daughter cells by attachment to host mitotic chromosomes, and this is thought to be mediated by the E2 protein. The actual partitioning strategy has been unclear and various models have been proposed, as shown in Figure 1. Early studies of BPV1 debated whether papillomavirus genome replication was licensed (each gnome is replicated once per cell cycle) or underwent random choice selection in the maintenance phase of replication (27, 28). Hoffman and colleagues showed that in different cell types, either mechanism could be used and expression of the E1 protein could promote genome amplification and random choice replication (11). Bienkowska-Haba and colleagues recently showed that HPV genomes were often lost to the cytosol in mitosis and copy number was maintained by S-Phase amplification (12).

In our study we observe clear ParB puncta in the nucleus of cells containing HPV18-ANCH genomes and clear attachment of these genomes to host chromosomes during mitosis. The puncta are heterogenous in size and intensity, suggesting that they contain more than one viral genome. In support of this, the HPV18-ANCH genomes are present at several hundred copies per cell, but much fewer puncta are observed (∼6-14 per cell). As shown in Figures 7 and 8, the genome-ParB-GFP puncta are partitioned to daughter cells in approximately, but not always exactly equal numbers. Furthermore, it is quite infrequent to observe faithful partitioning where a single punctum splits into two daughter punta as chromosomes separate in metaphase-anaphase (13.4% puncta observed). Therefore, we conclude that HPV18-ANCH genomes are partitioned somewhat equally to daughter cells by random attachment to host chromosomes, similar to the mode described for KSHV (4).

While the ANCHOR™ system allows visualization of replicating viral genomes, there are some considerations. We are not actually visualizing HPV18 genomes directly but through binding of the ParB-GFP protein. As the protein does not contain any localization sequences and is close to the nuclear pore exclusion limit, ParB-GFP is predominantly cytoplasmic but there are low amounts present in the nucleus that can cooperatively bind to the ANCH element in the HPV18 genome in a sliding mode that involves condensing the ANCH DNA with transient ParB bridges (29). Remarkably, the HPV18-ANCH genomes are maintained for many cell doublings with bound ParB-GFP, but we cannot rule out that the observed partitioning of viral DNA is influenced by the ParB protein.

We also observed (similar to findings by Bienkowska-Haba and colleagues with HPV16 and HPV31 genomes) that some HPV18-ANCH genomes were often lost to the cytosol in mitosis (12). Here, the loss of genomes was already established upon NEBD, indicative of failed tethering or displacement from host chromatin independent of spindle forces. This is surprising as it has long been thought that one of the advantages of tethering to host chromatin is to prevent cytoplasmic activation of innate immune sensors (30). The HPV18-ANCH cells have a high viral genome copy number per cell and it is possible that this does not represent the situation in the basal cells of a lesion where genomes are often present at undetectable levels. Alternatively, the low copy in basal cells might require the S-phase amplification described by Bienkowska-Haba to ensure that viral genomes are not lost (12).

## Materials and Methods

### Plasmids and Cloning

Plasmids were obtained from Lonza (pmaxGFP) and Invivogen (pSelect-mcs-puro). HPV18 cloned in pBR322 has been described previously (31), as well as the URR-deleted HPV18 genome (15) The URR-replicon has been described previously (15) as have HPV18 E1 and E2 expression vectors (32). The ParB-GFP and ANCH3 sequences were generated by PCR from an HPV18-ANCH-ParB-GFP genome obtained from the Schelhaas lab.

To generate URR-replicons with a polylinker at one of four cloning sites, a double-stranded oligonucleotide containing unique restriction sites was inserted into the NcoI, NheI, BlpI, or SbfI sites of the URR-replicon resulting in eight replicons with the polylinker in two orientations at each site. To clone the pmaxGFP URR-replicon, the GFP open reading frame was generated by PCR from pmaxGFP and inserted into the multiple cloning site of pSelect-mcs-puro (Invivogen) to generate pSelect-pmaxGFP. Similarly, the ParB-GFP ORF was PCR amplified and inserted into pSelect-mcs-puro. The resulting gene cassettes containing the hEF1/HTLV promoter and SV40 polyadenylation signal were PCR amplified and inserted into the URR-replicon. To generate HPV18-ANCH, the ANCH3 region was PCR amplified and inserted between two Asp718 sites in the late region of the HPV18 genome. ANCH3 is a specific bacterial chromosome partition sequence; the ANCHOR™ system is the property of NeoVirTech SAS (requests of use). URR-Replicons were transformed and grown in GT115 *E.coli* with 50 µg/mL Kanamycin LB-broth. The presence of multimerized URR-replicons in bacteria can be minimized by culturing them at 30°C instead of 37°C.

### HPV genome recircularization

HPV18 and HPV18-ANCH genomes were cleaved with EcoRI to release them from the pBR322 vector and recircularized with T4 DNA ligase at 5 µg/mL.

### Cell culture

Keratinocytes were cultured in Rheinwald-Green F medium (3:1 [v/v] F-12 [Ham]-DMEM, 5% FBS, 0.4 µg/mL hydrocortisone, 5 µg/mL insulin, 8.4 ng/mL cholera toxin, 10 ng/mL EGF, 24 µg/mL adenine, 100 U/mL penicillin and 100 µg/mL streptomycin) on a layer of irradiated J2-3T3 murine fibroblasts. Cells selected with G418 were co-cultured with G418-resistant J2-3T3 murine fibroblasts.

### Transient transfection and monitoring of fluorescent protein expression

Plasmids were transfected into both HPV18-negative and positive keratinocytes using Promega’s FuGENE® 6. Expression of florescent proteins was detected and quantified in live cells using an IncuCyte SX5 (Sartorius).

### Electroporation of Human foreskin keratinocytes (HFKs)

HFKs were cultured with 10µM Y-27632 and electroporated with recircularized HPV18 or HPV18-ANCH genomes using the Amaxa^TM^ Human Keratinocyte Nucleofector^TM^ Kit (Lonza). Following transfection, cells were plated on irradiated fibroblasts for downstream assays (replication, colony establishment assay and long-term passage). Y-27632 was omitted from the medium one-day post-transfection. URR-replicons were electroporated similarly into the established HPV18 or HPV18-ANCH cell lines.

### Colony Establishment Assay

Following electroporation, 2 × 10^3^ cells were plated on irradiated fibroblasts for replicon establishment colony assays. Cells were allowed to recover for one-day and then selected with 400 µg/mL G418 for four days, and 200 µg/mL for approximately two weeks. Colonies were fixed with 3.7% formaldehyde in PBS and stained with 0.14% methylene blue.

### Southern blotting

Total DNA was extracted from cells using the DNeasy Blood and Tissue kit (Qiagen). For transient replication assays, DNA was harvested five to six days post-transfection without selection. For stable replication analysis, DNA was extracted at least at pass four or greater after G418 selection. For transient replicon analysis, 2 µg DNA was digested with AflII and DpnI (to digest unreplicated DNA) or BstXI and DpnI. DNA was separated on a 1% agarose gel. For stable HPV18 genome analysis, 3 µg DNA was digested with XhoI or PacI to cleave cellular DNA (HPV18 non-cutters), or AflII to linearize the genome and separated the bands on agarose gel. DNA was transferred to nylon membranes using the TurboBlotter downward transfer system (Whatman). Following DNA transfer, membranes were UV-crosslinked, dried, and hybridized with 25ng ^32^P-labeled probe in 3X SSC, 2% SDS, 5X Denhardt’s, 200µg/ml salmon sperm DNA. To detect replicons, radioactive probes were generated with ^32^P-dCTP-labeled URR-negative replicon to avoid cross-hybridization with the viral genome. For HPV18-ANCH analysis, ^32^P-labeled HPV18-ANCH DNA was used as a probe. Radiolabeled probes were generated using a Random Prime DNA labeling kit (Roche). Membranes were washed in 0.1% SDS/0.1 X SSC and ^32^P-hybridized DNA was visualized and quantitated with a Typhoon phosphorimager (GE Bioscience).

### Combined Immunofluorescence and Fluorescent *In Situ* Hybridization (FISH)

Cells were fixed in 4% PFA in PBS at room temperature for 15 minutes, permeabilized, and blocked, and stained with an anti-CometGFP (also recognizes Dasher) antibody (ATUM). After immunostaining, cells were fixed at room temperature with methanol and acetic acid solution (3:1 v/v) for 10 minutes and 2% PFA for 1 minute. Coverslips were treated with an RNace-iT cocktail and dehydrated in a 70%, 90%, and 100% ethanol series and air dried. DNA FISH probes were prepared using the FISH-Tag DNA Multicolor Kit labeling kit (Life Technologies). Hybridization was performed overnight in 1X Hybridization Buffer (Empire Genomics) with 50-75ng of labeled probe DNA at 37°C. Slides were washed at room temperature with 1X phosphate-buffered detergent (PBD, MP Biosciences), followed by washing with wash buffer (0.5X SSC, 0.1% SDS) at 65°C. Nuclei were stained with DAPI and coverslips were mounted using Prolong Gold (Life Technologies).

### Confocal microscopy

Images were collected with a Leica SP8 WLL TAN 405 HyD DMi8 AFC laser scanning confocal microscope and processed using Leica LAS AF Lite Software (Leica Microsystems).

### Live cell imaging

Keratinocytes were co-cultured with neomycin resistant irradiated fibroblasts on chambered µ-slides (Ibidi) and imaged in a Tokai Hit 5% CO_2_ chamber at 37°C for up to several days using a Leica SP8 confocal microscope with or without live-cell dyes (cytoskeleton, Inc). Cells treated with live-cell dyes were given pre-warmed F-medium with 50 μM/mL G418 containing 1X SPY-650 (DNA) and 1X SPY-555 (tubulin) dyes and incubated at 37°C 5% CO_2_ for 1hr. After 1hr, media was replaced with F-medium 50 μM/mL G418. Confocal images were collected using a Leica TCS-SP8 laser scanning confocal microscope equipped with a HC PL APO 63X/1.4OIL CS2 objective (Leica Microsystems) with a 200µm x 200µm x 15µm field of view, 1µm z-step size, 1 airy unit (AU) optical sections. Time-lapse images recorded with 13-minute time intervals for 12-72 hours.

### Image analysis

For Figures 4-6, images were processed using Leica LAS AF Lite software (Leica Microsystems), for Figure 9A, images were MAX intensity projected in the z plane and denoised with Gaussian blurring (sigma=0.5) in Fiji/ImageJ (33). Mitosis stages were defined by DNA morphology (34). The NEBD event and the post-mitotic G1 stage were detected by the influx and efflux, respectively, of a non-specific cytosolic signal from ParB-GFP into the nuclear volume (35). Cells from two independent experiments (n=7 and n=5) with low ParB-GFP expression, which progress through mitosis normally, were manually selected and classified into mitotic stages based on chromatin and cytosol signals. The two nascent daughter cells were quantified as one entity. Image quantification was performed using Imaris v10.2 (Oxford Instruments). Chromatin was segmented using the “surfaces” function and filtered using the “absolute intensity threshold”, “volume”, and “ellipticity” thresholds. Punctae were segmented with the ‘Spots’ function (spot size 1.5 um), filtered by ‘Intensity centre threshold’, punctae outside the target cell were manually deleted. Puncti were then classified with ‘spots classification’ tool by partial or complete localization within ‘chromatin’ surface and the rest classified as ‘cytosol’. Statistical analysis Kolmogorov-Smirnov test (36) and two tailed t-test (37) and plotting of quantified punctae were performed using Microsoft Office Excel (Microsoft). Plots and figure assembly were done in Affinity designer (Serif Europe).

## Acknowledgments

This work was funded by the Intramural Research Program (Division of Intramural Research, DIR), National Institute of Allergy and Infectious Diseases (ZIA AI001073 to AAM) and the European Research Council (ERC consolidator grant 682899 MitoVIn to MS). We thank JJ Miranda and members of the McBride laboratory for feedback on the manuscript.

---

Supplemental Figure 1. Optimization of the URR-Replicon Expression Vector

A polylinker was inserted into four positions in the URR-replicon (in the sense and antisense orientation) between functional elements. Thereafter, each plasmid was electroporated into HFK, or HFK-HPV18 cells to assess their ability to establish as stable URR-replicons. This resulted in stable neomycin-resistant colonies for all plasmids, but only in the presence of HPV18. pCGneo, a neomycin resistance plasmid, does not contain the HPV18 URR and therefore cannot replicate. NB: Blp sense has Increased colonies due to seeding error.

Supplemental Figure 2. Developing an optimal expression cassette

(A) fluorescent protein gene expression cassettes pmaxGFP and pSelect-pmaxGFP. The plasmids shown were transfected into HPV negative or HPV18 containing HFKs and imaged in a IncuCyte scanner. (B) Total integrated fluorescent intensity at 48 hours post-transfection (n=3, error bars=SD). (C) The green cell count at 48 hours post-transfection (n=3, error bars=SD).

Supplemental Figure 3. Replication of the pmaxGFP-URR-replicons in primary HFKs.

(A) Map of the URR replicon and cloning positions for pMAX-GFP cassettes. (B) Southern blot assay. The eight pmaxGFP-URR-replicons and pCGneo (pNeo) plasmid were electroporated into HFK-HPV18 cells. DNA was harvested after six days; the cellular DNA in the top panel was digested with AflII and in the middle panel with AflII and DpnI to linearize and detect DpnI resistant replicated DNA. Size/copy number markers are loaded on the left. The top and middle membranes were probed with a URR-deleted replicon, and the bottom with the HPV18 genome. The endogenous HPV18 genomes were maintained at several-hundred copies per cell. (C) The transfected cells were grown with G418 selection for ∼4 weeks and genomic DNA isolated The top panel shows a Southern blot with DNA cleaved with AflII to linearize the replicon, and the bottom panel with PacI (a non-cutter) to determine the extra-chromosomal status of the endogenous HPV18 genome. Probing the top panel with pMAX-GFP showed that, while most replicons were present, they had undergone significant rearrangements and deletions. The bottom panel (probed with the HPV18 genome) shows that the HPV18 genomes remained extra-chromosomal and at high copy number after ∼4 weeks.

Supplemental Movie 4. HPV18 genomes visualized in dividing cells by ANCHOR technology.

Cell line HFK HPV18-ANCH a/s ParB-GFP-pool (Table 1) HPV18 genomes/ParB-GFP are in green, host DNA is blue (SPY-650 DNA dye) and microtubules are in red. Time-lapsed image collections were deconvolved using Huygens Professional compute engine 24.10.0p2 64b, and visualized and recorded in 3D View in Bitplane Imaris v10.2.0.

## Notes

### Competing Interest Statement

The authors have declared no competing interest.

