## Supplementary Figures 1-3 for "Tracking Replicating HPV Genomes in Proliferating Keratinocytes"

Supplemental figure 1

A

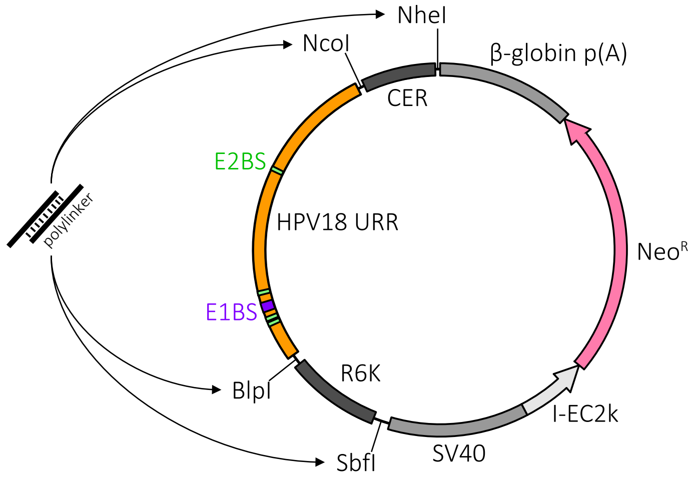


B

Polylinker sequence

| SacII | XbaI | BsiWI | EcoRV | ClaI | BstBI |
|---|---|---|---|---|---|
| CCGC^GG | T^CTAGA | C^GTACG | GAT^ATC | AT^CGAT | TT^CGAA |
| GG^CGCC | AGATC^T | GCTAG^C | CTA^TAG | TAGC^TA | AAGC^TT |

C

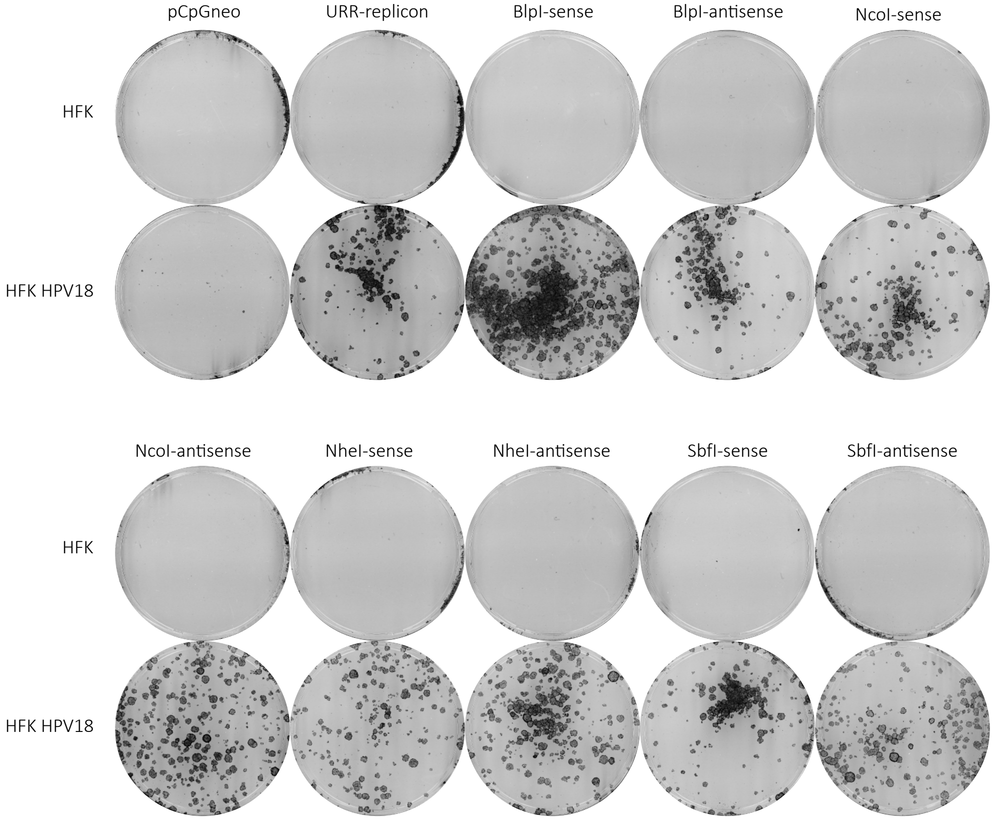

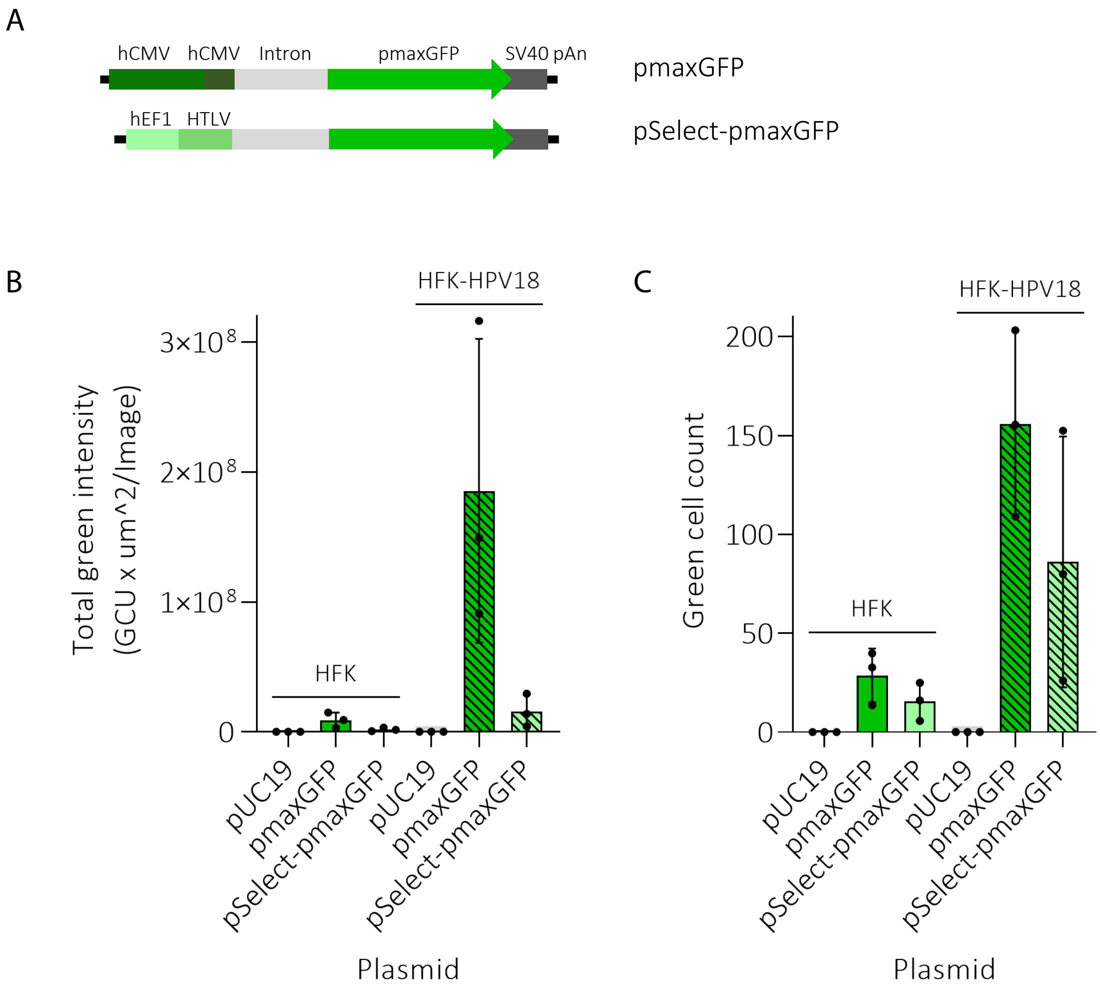
A
hCMV
hCMV
Intron
pmaxGFP
SV40 pAn
pmaxGFP
hEF1
HTLV
pSelect-pmaxGFP
B
HFK-HPV18
Total green intensity (GCU x um^2/Image)
3×10^8
2×10^8
1×10^8
0
HFK
pUC19
pmaxGFP
pSelect-pmaxGFP
pUC19
pmaxGFP
pSelect-pmaxGFP
Plasmid
C
HFK-HPV18
Green cell count
200
150
100
50
0
HFK
pUC19
pmaxGFP
pSelect-pmaxGFP
pUC19
pmaxGFP
pSelect-pmaxGFP
Plasmid

Supplemental Figure 3

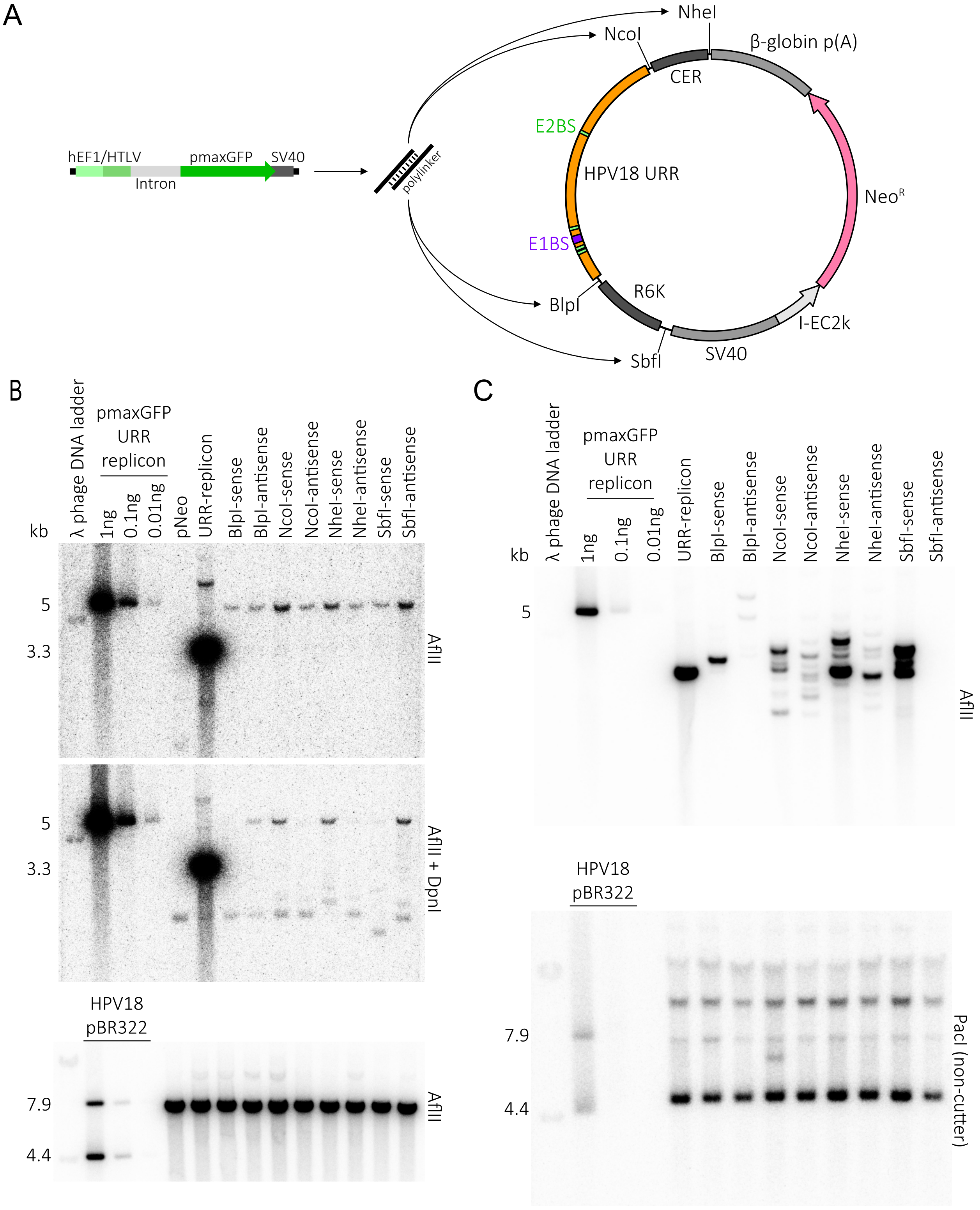
